# A comprehensive atlas of fetal splicing patterns in the brain of adult myotonic dystrophy type 1 patients

**DOI:** 10.1101/2021.10.01.462715

**Authors:** Max J. F. Degener, Remco T.P. van Cruchten, Brittney A. Otero, Eric T. Wang, Derick G. Wansink, Peter A.C. ‘t Hoen

## Abstract

In patients with myotonic dystrophy type 1 (DM1), dysregulation of RNA-binding proteins like MBNL and CELF1 leads to alternative splicing of exons and is thought to induce a return to fetal splicing patterns in adult tissues, including the central nervous system (CNS). To comprehensively evaluate this, we created an atlas of developmentally regulated splicing patterns in the frontal cortex of healthy individuals and DM1 patients by combining RNA-seq data from BrainSpan, GTEx and DM1 patients. Thirty four splice events displayed an inclusion pattern in DM1 patients that is typical for the fetal situation in healthy individuals. The regulation of DM1-relevant splicing patterns could partly be explained by changes in mRNA expression of the splice regulators *MBNL1, MBNL2* and *CELF1*. On the contrary, interindividual differences in splicing patterns between healthy adults could not be explained by differential expression of these splice regulators. Our findings lend transcriptome-wide evidence to the previously noted shift to fetal splicing patterns in the adult DM1 brain as a consequence of an imbalance in antagonistic MBNL and CELF1 activities. Our atlas serves as a solid foundation for further study and understanding of the cognitive phenotype in patients.

## INTRODUCTION

Myotonic dystrophy type 1 (DM1; OMIM #160900), also known as Steinert’s disease or dystrophia myotonica, is an autosomal dominant neuromuscular disease with a highly variable, multisystemic clinical presentation, affecting skeletal and smooth muscles, the central nervous system (CNS), the heart and several other organs. DM1 is the most common form of adult onset muscular dystrophy with an estimated global prevalence of 1:8,000 (1). The clinical phenotype of DM1 is most notably defined by myotonia, a delayed relaxation of skeletal muscles following contraction. Additionally, affected individuals show a variable combination of progressive weakness of distal muscle groups, insulin resistance, cardiac arrhythmia, cataract, fatigue, cognitive impairment and changes in personality and behaviour.

The cause of DM1 is a (CTG)n trinucleotide repeat expansion in the 3’-untranslated region of the DM1 protein kinase (DMPK) gene, located on the long arm of chromosome 19 (2). The central mechanism of DM1 pathophysiology involves the formation of RNA hairpin structures by the (CUG)n repeat in DMPK transcripts (3, 4). Proteins of the muscleblind like splicing regulator (MBNL) family are recruited by these hairpin structures, bind to repetitive “YGCY” motifs of the (CUG)n expansion and form RNA foci (5–7). Consequently, MBNL proteins, trapped in nuclear RNA foci, are depleted from their normal RNA targets (8). Since MBNL proteins also regulate their own splicing, the depletion of MBNL proteins leads to a further loss of MBNL function due to the formation of other splice variants (9). Furthermore, the presence of expanded DMPK transcripts increases levels of hyperphosphorylated, stable CUG-binding protein and ETR3-like factor 1 (CELF1) (10).

RNA-binding proteins (RBPs) like CELF1, MBNL1 and MBNL2 function as trans-acting regulators of alternative splicing (11, 12). These RBPs regulate alternative splicing by binding to short RNA motifs in the pre-mRNA. Depending on the binding sites and the RBPs involved, splicing of the target RNA elements can be promoted or suppressed (13). Although the functional impact of many individual splice events remains ill understood, the coordination of these events, driven by binding of RBPs, determines the development of tissues, particularly of the heart, skeletal muscle and brain (14). During heart development in mice and chickens, MBNL1 and CELF1 mediate a highly conserved transition from fetal to adult splicing patterns (15). Similarly, loss of Mbnl1 and Mbnl2 in the brain of adult knockout (KO) mice shifts the splicing profile to an earlier stage of CNS development and causes mis-splicing of several important developmentally regulated exons (16, 17).

A cornerstone of DM1 pathophysiology is the aberrant alternative splicing of many pre-mRNA products and the preferential expression of fetal transcript variants in the adult state. Numerous studies revealed mis-splicing in DM1 patient tissue and animal models by means of RT-PCR, microarray and RNA-Sequencing (e.g. (17–24)) Mis-splicing in DM1 is not limited to one type of tissue, but affects virtually all systems associated with the disease, in particular skeletal and cardiac muscles as well as the CNS (19, 20, 22). Importantly, abnormal splicing of certain genes has been linked to characteristic DM1 symptoms, albeit with variable levels of evidence (17, 19, 25–27). The alternative splicing of a number of these phenotype-linked genes is regulated by MBNL1 or MBNL2 (DMD, BIN1, SCN5A), CELF1 (CLCN1, RYR1, INSR, TNNT2) or both (CACNA1S, PKM, GRIN1, MAPT). Therefore, the loss-of-function of MBNL1/2, due to their sequestration in foci, and the gain-of-function of CELF1, due to its hyperphosphorylation, could explain the dysregulation of alternative splicing in the disease state (7, 28). MBNL1/2 and CELF1 regulate splicing in an antagonistic fashion by either promoting adult (MBNL1/2) or fetal (CELF1) splicing patterns (29, 30). Consequently, the altered activity of MBNL1/2 and CELF1 will induce the expression of fetal transcript variants in adult DM1 tissues (8) and DM1 mouse models (24).

Here, we set out to create an atlas of all developmentally regulated splicing patterns in the human frontal cortex and studied to what extent they were affected in the brains of DM1 patients. For this, we analyzed publicly available RNA-Seq data from healthy individuals, generated by the BrainSpan and GTEx projects (31, 32), together with recently published data from the brains of DM1 patients and controls (23). In contrast to previous more targeted approaches, this now allowed us to make a comprehensive inventory of splicing events demonstrating a transition towards the fetal state in the DM1 frontal cortex. Moreover, we assessed to what extent this could be explained by the RNA expression levels of CELF1 and MBNL1/2.

## METHODS

### Datasets

The BrainSpan Atlas of the Developing and Adult Human Brain is a resource for studying the spatial and temporal development of the human brain (31). The RNA-Seq dataset consists of multiple CNS regions and age groups (from eight weeks before birth to 40 years of age). The BrainSpan consortium only included samples that did not have any confounding pathological factors (e.g. cerebrovascular incidents, tumors), did not show chromosomal abnormalities or malformations and lesions, had not been subjected to drug or alcohol abuse and showed a RNA integrity number (RIN) of at least 5. Our selection consisted of 139 samples from 39 individuals (23 male, 16 female) in which four subregions of the frontal cortex were represented (i.e. dorsolateral: N=35, ventrolateral: N=36, medial: N=35 and orbital: N=33) (Figure 1A). Since we found no differences in splicing of our exons of interest between these subregions, this data was pooled in further analyses (Supplemental Figure S1, Supplemental Table 1). The RNA-Seq was paired-end with a read length of 76 bp. Access to FASTQ files and comprehensive phenotype data was requested from the database of Genotypes and Phenotypes (dbGaP; study accession: phs000731.v2.p1). FASTQ files were downloaded from the sequence read archive (SRA).

**Figure 1:**
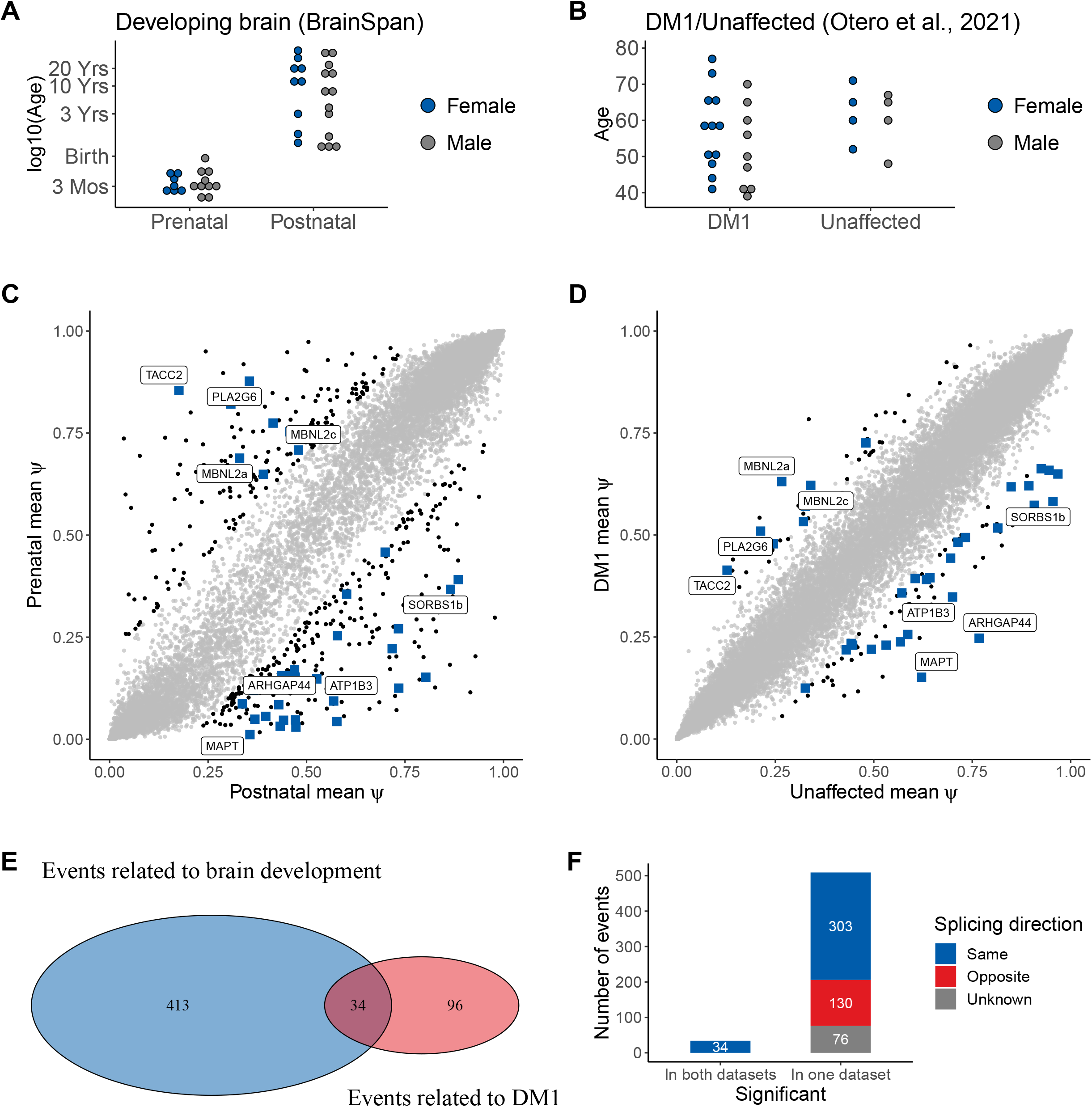
A quarter of the aberrant splice events in adult DM1 brains are associated with brain development. A) Overview of the developmental stage of 139 Brainspan RNA-Seq samples from frontal cortices of 39 healthy donors in the developmental dataset. B) Overview of the DM1 dataset from Otero et al. (2021) containing RNA-Seq samples from the frontal cortices of 21 DM1 adult patients and 8 unaffected controls. C) Scatter plot of Ψ in prenatal (y-axis) and postnatal (x-axis) samples for all measured splice events in the developmental dataset. The 447 splice events with significant differences between groups (|ΔΨ| > 0.2, p < 0.01 by rank-sum test) are in black. Events shaped as a blue-colored square were significantly different in both datasets and those with the largest mean ΔΨ in the DM1 dataset are labeled by their name (see also Figure 2). D) Scatter plot of mean Ψ in DM1 (y-axis) and healthy (x-axis) samples for all splice events measured in the DM1 dataset. The 130 splice events with significant differences between groups (|ΔΨ| > 0.2, p < 0.01 by rank-sum test) are in black. Further labeling also as in C. E) Venn diagram of splice events that featured a marked and significantly different change (|ΔΨ| > 0.2, p < 0.01 by rank-sum test) between prenatal and postnatal samples (i.e. related to brain development), and between DM1 and unaffected samples (i.e. related to DM1). F) Histogram of splice events with similar or opposite direction for the events significantly changed in both the Brainspan and the DM1 datasets (left bar) of in just one of the datasets (right bar). Here, events were considered significant when p < 0.01 by rank-sum test in both datasets. The 34 high-confidence events all show the same direction of change in their exon inclusion.

The Otero et al. (2021) dataset consists of frontal cortex samples from 21 DM1 patients (9 male, 12 female) and 8 controls (4 male, 4 female) (Figure 1B, (23)). The samples were retrieved from frozen brain tissue of adult individuals. The age of the donors ranged from 39 to 77 years (median = 59). Controls were labelled as “unaffected”. The RNA-Seq was paired-end with a read length of 75 bp. More details can be found in the associated publication (23). The raw RNA-Seq data were downloaded from GEO (study accession: GSE157428).

The GTEx project provides RNA-Seq data from a variety of tissues from healthy donors (32). We focused on the human frontal cortex and included only samples with RIN above 7, coming from donors with non-disease related death (Hardy Scale below 3), in which autolysis of the extracted tissue was not visible upon inspection by a pathologist. The final selection consisted of 54 samples from 54 donors (42 male, 12 female). The age of the donors ranged from 23 to 70 (median = 60.5). The RNA-Seq was paired-end with a read length of 76 bp. For the selected samples, raw gene-level counts were downloaded from the public GTEx portal (release V8). Access to BAM files and comprehensive sample and phenotype data was requested from dbGaP (study accession: phs000424.v8.p2). Following approval, selected BAM files and metadata were downloaded from the corresponding repository on the NHGRI AnVIL (Genomic Data Science Analysis, Visualization, and Informatics Lab-space) platform.

### RNA-Seq data processing

To allow comparison across datasets, we applied the same RNA-Seq pre-processing and expression quantification steps on FASTA files from BrainSpan and Otero et al. (2021) as specified for GTEx V8 (https://gtexportal.org/home/documentationPage). FASTQ files were aligned to the hg38 genome with STAR 2.7.0f (33). The STAR index was built with the genome sequence (GRCh38.p10) in FASTA format and the comprehensive gene annotation, both retrieved from GENCODE release v26. Samtools 1.9 was used to sort and index the resulting BAM files (34). Gene-level counts were calculated by RNASeQC 2.3.5 (35) and normalized to log2 counts per million (CPM) with the trimmed mean of M-values method, as implemented by the edgeR 3.28.0 package (36). RNASeQC 2.3.5 required a collapsed annotation file which lists only a single transcript per gene (35). The collapse_annotation.py script from GTEx was used to collapse the comprehensive GENCODE v26 gene annotation (https://github.com/broadinstitute/gtex-pipeline/tree/master/gene_model). Pathway analysis with the May 2021 release of Wikipathways (37) was performed using ClueGO v2.5.8 (38) with GO term fusion and the Bonferroni right-sided post test.

### Calculation of Percent Spliced-In

To estimate exon inclusion for exon skipping events, percent spliced-in (PSI, Ψ) values were computed with MISO 0.5.4 (39). MISO estimates the Ψ for a given splice event by using Bayesian inference on a combination of inclusion reads, exclusion reads, junction-spanning reads and upstream/downstream exonic reads. Splice events which were not detected in each sample within a dataset were excluded from that dataset. The function gff_make_annotation of rnaseqlib 0.1 (40) was used to generate MISO annotation files of the most recent genome build (hg38) based on the comprehensive GENCODE v26 annotation in the UCSC genePred format (http://hgdownload.soe.ucsc.edu/goldenPath/hg38/database/wgEncodeGencodeCompV26.txt.gz).

### Calculation of correlations and partial correlations

Ordinal relationships between estimates of exon inclusion and RNA expression were assessed by computing the Spearman’s rank correlation. This measure of correlation was chosen in favor of the Pearson correlation since the Ψs for the majority of splice events were skewed and not normally distributed, as assessed by the Shapiro-Wilk test. The function corr.test of the psych 1.8.12 R package was used to compute the correlation coefficients, to test for significant correlations and to control the false discovery rate (FDR) at 5% with the Benjamini-Hochberg procedure (41, 42). Correlations were assessed (1) between Ψ estimates of selected splice events and age of the sample donor and (2) between Ψ estimates and RNA expression of selected RBPs (i.e. CELF1, MBNL1, MBNL2) and (3) between Ψ estimates of all high-confidence splice event pairs (i.e. pairwise cross-correlations). To account for the influence of other variables on the pairwise correlations between splice events, partial correlations were computed with the ppcor 1.1 R package (43). A partial correlation represents the association of two variables (e.g. inclusion of exon A and exon B) while removing the effect of one or more other variables (e.g. RNA expression of gene A and/or gene B). The age of the sample donor, the disease state (DM1 or unaffected) and the RNA expression level of the RBPs were separately taken into account for the partial correlation analysis. The resulting controlled partial correlations were compared to the zero-order pairwise correlations.

## RESULTS

### Splicing patterns in the frontal cortex of adult DM1 patients resemble those of developing brains

DM1-related aberrant splicing has been associated with the expression of fetal transcript variants (17, 26, 44–47). We performed a comprehensive evaluation of the developmental regulation of DM1-related exon skipping events by comparing exon inclusion in frontal cortex samples from the human BrainSpan dataset (Figure 1A) and from a recent dataset including 21 DM1 patients and 8 controls (Figure 1B) (23, 31). PSI (Ψ) values were calculated for all splice events and reflect the inclusion rate of a cassette exon that is either included or skipped within the boundaries of its flanking upstream and downstream exons.

In the developing brain dataset, we identified 447 events with significant differences (|ΔΨ| > 0.2, p < 0.01 by rank-sum test) between prenatal and postnatal frontal cortex (Figure 1C; Supplemental Table 2). One hundred thirty events displayed significant differences (|ΔΨ| > 0.2, p < 0.01 by rank-sum test) between DM1 and healthy frontal cortex samples (Figure 1D; Supplemental Table 3; (23)). A significant overlap of 34 exons (p < 2.2 e-16, Fisher’s exact test) was found between the sets of splice events in the developing brains and the DM1 brains (Figure 1E; Supplemental Table 4). These 34 events were present in 30 different genes and involved 32 unique cassette exons. When there were multiple events in the same gene, these events were labelled with a/b/c etc. (see Supplemental Table 4 for the exact nomenclature). Two cassette exons were present in multiple events differing by a variable 5’ end (SORBS1a, SORBS1b) or the upstream exons (MBNL2b, MBNL2c). Compared to the background of all differentially spliced genes in the DM1 dataset, these 30 were enriched (Bonferroni adj. p = 0.025) for genes in the Wikipathway ‘ectoderm differentiation’, represented by DMD, KIFC3, NUMA1 and TCF3 (37). Notably, all 34 splice events changed towards the prenatal inclusion pattern in the DM1 brains (Figure 1F) and the absolute ΔΨ for DM1 versus control was on average larger in this subset of 34 events compared to all 130 events related to DM1 (p < 0.01 by rank-sum test, Supplemental Figure S2). Together this confirms, on a transcriptome-wide scale, previous observations that DM1 brains reflect embryonic / fetal splicing patterns. The list of 34 splice events contains many of the previously described aberrantly spliced exons in DM1 and known MBNL targets, such as MBNL1 exon 5, MBNL2 exon 5, DMD exon 78, and MAPT exon 3.

Closer inspection of the developmental regulation of splicing for these 34 splice events demonstrated gradual changes from the fetal through the perinatal to the postnatal period. In Figure 2, we display the splice events with the largest mean difference in Ψ between the DM1 frontal cortex and that of controls. For all splice events with an increase of Ψ during development, Ψ values were lower in DM1 patients than in controls, and vice versa. The strongest increases in Ψ throughout development were observed in ARHGAP44, ATP1B3, MAPT and SORBS1 transcripts, whilst the strongest decreases in Ψ were observed in MBNL2, PLA2G6 and TACC2. For an overview of Ψs of these exons across CNS subregions, see Supplemental Figure S3. For most of these genes, pronounced changes in splicing patterns were observed during the perinatal period, but the timing and the extent to which these patterns changed from the perinatal period to the adult stage differed considerably between exons.

**Figure 2:**
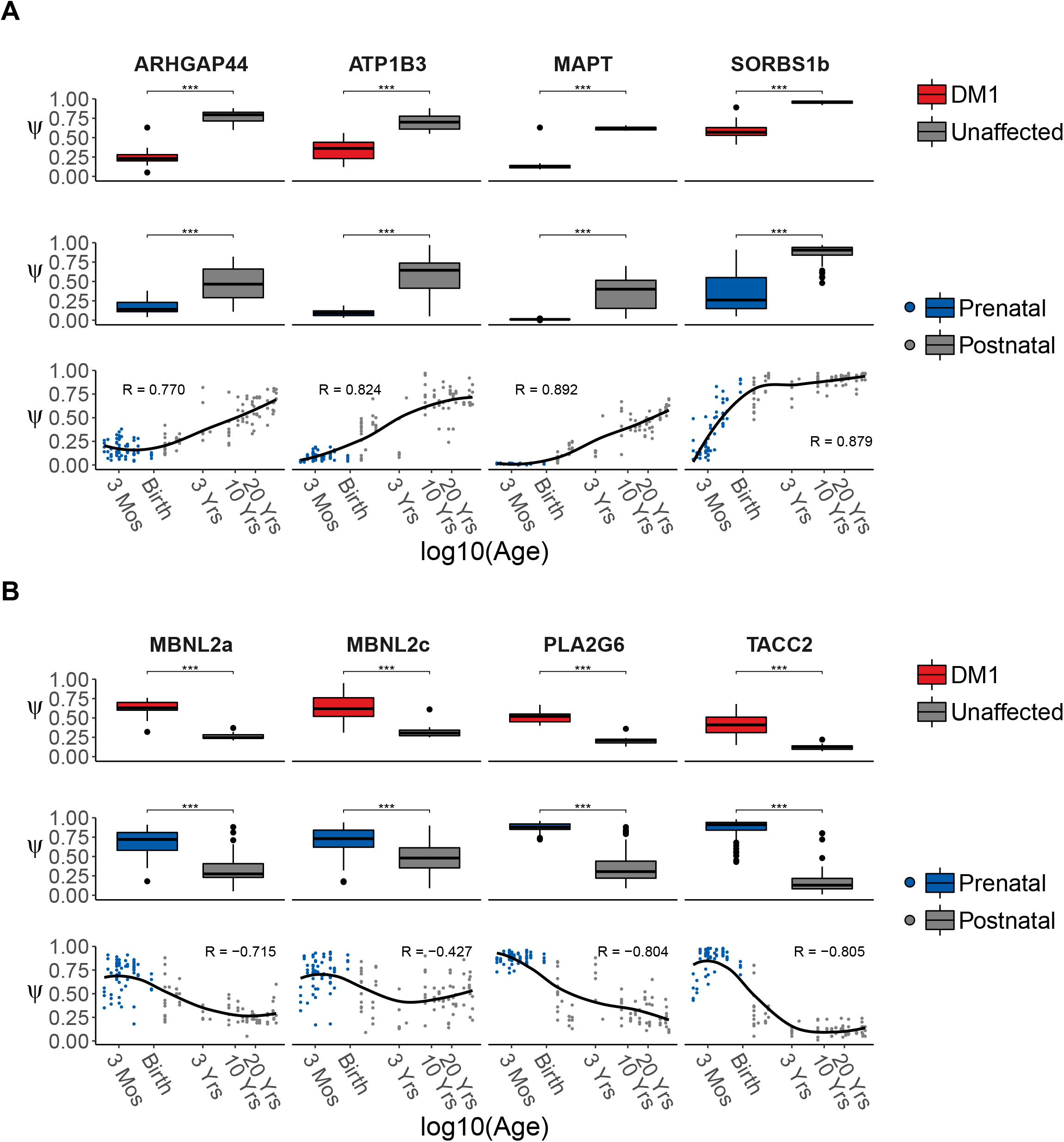
Splice events with strongest up- and downregulation in DM1 show gradual change through development. Splice events with the largest decrease (A) or increase (B) in Ψ in DM1 patients compared to unaffected adults. The top rows in panels A and B show boxplots of Ψs for events selected for the largest difference in mean Ψ between frontal cortex samples from DM1 patients and unaffected adults. The boxplots in the middle rows are based on frontal cortex samples from healthy prenatal and postnatal donors. A significant difference between groups was assessed by the rank-sum test. P-values were FDR-corrected with the Benjamini-Hochberg procedure (ns: p > 0.05,*: p < 0.05,**: p < 0.01,***: p < 0.001). The bottom row shows scatterplots of Ψ for the selected events in the healthy, developing human frontal cortex. Post-conceptional age was recorded in days and plotted on a log10 scale. Arbitrary x-axis labels were chosen to highlight specific timepoints during development (Mos=Months, Yrs=Years). The regression curve was estimated by the LOESS method.

### Expression of *MBNL* and *CELF1* in the developing and DM1 brain

Since RBPs are known to regulate alternative splicing during human tissue development, we inspected the RNA expression levels of RBPs that have been linked to DM1 pathophysiology, i.e. those of the MBNL and CELF families, throughout development and in the disease situation (7, 14). For CELF1, as well as CELF2, -3, -5 and -6, we observed a significant (FDR < 0.05) downregulation in prenatal and postnatal samples (Figure 3 and Supplemental Figure S4; log2 fold change for CELF1 (logFC_CELF1_) = -1.23). On the contrary, for MBNL1 and MBNL2, as well as CELF4, expression increased gradually and significantly (FDR < 0.05; logFC_MBNL1_ = 1.60; logFC_MBNL2_ = 2.66) during development, whereas no change was observed for MBNL3 (Supplemental Figure S5). The RNA expression levels of CELF1, -3 and -6 did not differ between DM1 cases and controls, while a slight upregulation of MBNL1, 2 and -3 was found in DM1 (logFC_CELF1_ = -0.09; logFC_MBNL1_ = -0.50; logFC_MBNL2_ = -0.44, logFC_MBNL3_ = -0.60). For CELF2, -4 and -5 we noticed a subtle downregulation in DM1 (Supplemental Figure S4). Since MBNL1/2 and CELF1 are best characterized in the context of DM1, and most of their family member featured similar or only modestly different effects, we focused on CELF1 and MBNL1/2 for further analyses.

**Figure 3:**
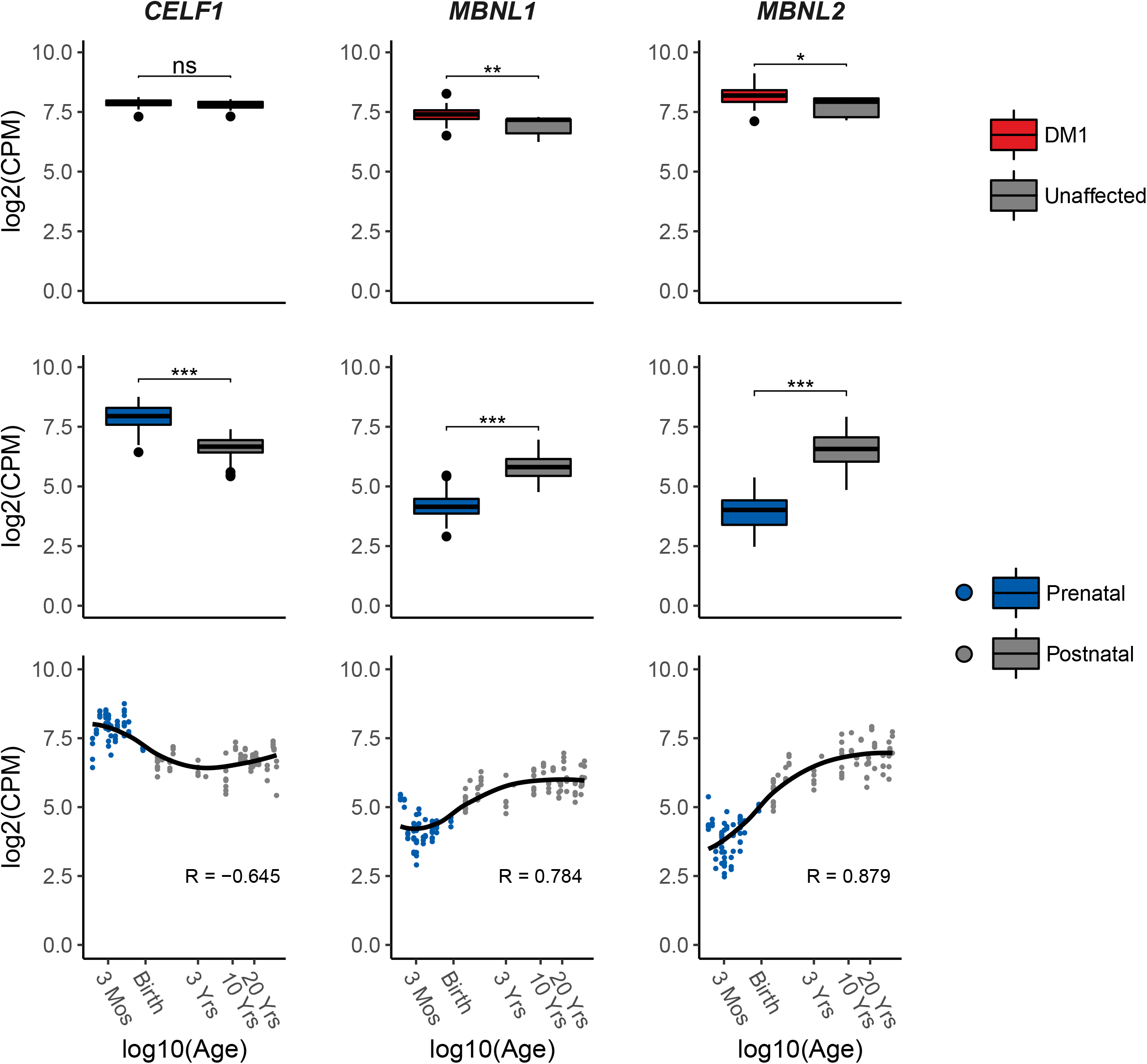
Differences in RNA expression of DM1-relevant splicing factors throughout human brain development and between DM1 patients and unaffected adults. The top row shows boxplots of RNA expression (log2 of CPM) of *CELF1, MBNL1 and MBNL2* in frontal cortex samples from DM1 patients and unaffected adults. The boxplots in the middle row are based on frontal cortex samples from healthy prenatal and postnatal donors. Significant differences between groups were assessed by the rank-sum test. P-values were FDR-corrected with the Benjamini-Hochberg procedure (ns: p > 0.05,*: p < 0.05,**: p < 0.01,***: p < 0.001). The bottom row shows scatterplots of RNA expression (log2 CPM) of *CELF1, MBNL1 and MBNL2* in the healthy, developing frontal cortex. Post-conceptional age was plotted on a log10 scale. X-axis labels were chosen to highlight arbitrary timepoints during development (Mos=Months, Yrs=Years). The regression curve was estimated by the LOESS method.

### DM1-relevant developmental splice events are associated with expression of *CELF1* and *MBNL1/2*

Next, we assessed the potential effects of the balance between CELF1 and MBNL expression on the splicing patterns of the set of 34 DM1-relevant developmental splice events. To achieve this, we calculated correlations between CELF1 and MBNL expression on the one hand, and the Ψ of these splice events on the other hand. In the developing brain, we observed that all 8 splice events with increased inclusion in the prenatal state and in DM1 patients were strongly positively correlated with the expression of CELF1 and strongly negatively correlated with the expression of MBNL1 and MBNL2 (Figure 4 and Supplemental Figure S6, examples in Supplemental Figure S7A). Conversely, the 26 events with increased inclusion in the postnatal state and decreased inclusion in DM1 patients, were strongly negatively correlated with the expression of CELF1 and strongly positively regulated with the expression of MBNL1 and MBNL2. Such correlations were much less strong within the sets of prenatal or postnatal samples, indicating that changes during development are driving these correlations (Supplemental Figure S8). Collectively, these findings indicate a high level of coordination within this set of splice events during brain development, and suggest an association with the balance between the activity of the CELF1 and MBNL splicing factors.

**Figure 4:**
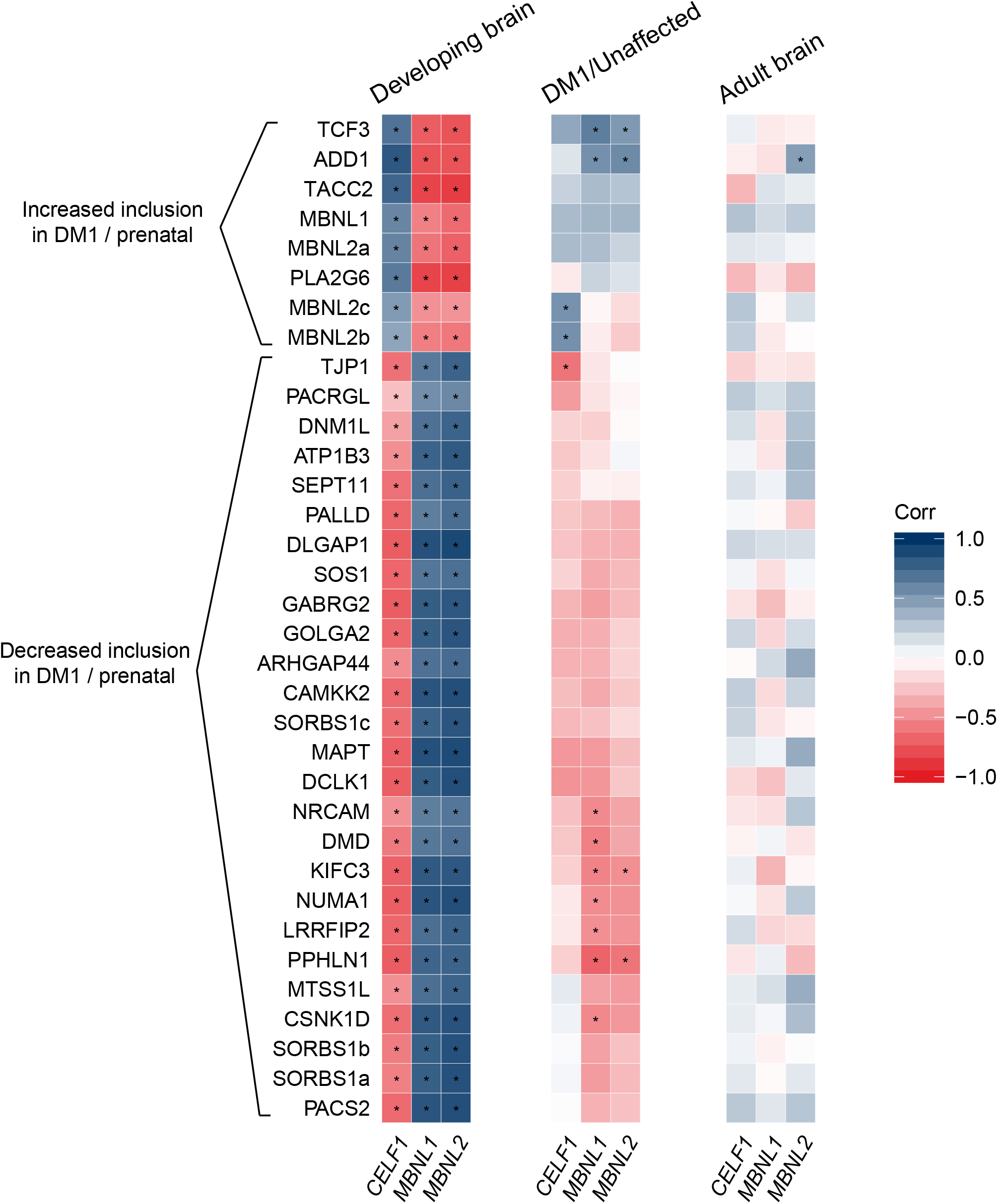
Correlation of alternative splicing and RNA expression of CELF1, MBNL1 and MBNL2. Correlation between Ψ of high-confidence exon skipping events and RNA expression levels of *CELF1, MBNL1* and *MBNL2*. Correlations are displayed for frontal cortex samples from the healthy, developing frontal cortex (left), from DM1 patients and unaffected adults (middle) and from the healthy, adult frontal cortex (right). The splice events are hierarchically clustered based on the average distance between the correlations for the DM1/unaffected samples. The color scale on the right reflects the value of the Spearman’s rank correlation coefficient. Asterisks indicate a significant correlation (per study FDR-corrected p < 0.05).

In the dataset of DM1 and control adult frontal cortex samples, we did not observe similarly strong correlations between the inclusion rates of the 34 DM1-relevant developmental splice events and the expression levels of CELF1 and MBNL (Figure 4 and Supplemental Figure S6, examples in Supplemental Figure S7B). However, upon closer inspection of these correlations, we can appreciate that the set of 8 splice events with increased inclusion in the prenatal state and in DM1 patients were positively correlated with the RNA expression of CELF1, MBNL1, and MBNL2. Again, the 26 events with increased inclusion in the postnatal state and decreased inclusion in DM1 showed the opposite pattern. The correlations between MBNL1/2 expression and splicing in the DM1 and unaffected brain showed opposite trends compared to the correlations observed in the developing brain. These trends were not observed when analyzing the DM1 and unaffected brain separately (Supplemental Figure S9).

Since the DM1 dataset contained only 8 control individuals, we performed a similar analysis in the much larger set of GTEx frontal cortex samples from healthy adult donors, which has a higher statistical power (N=54, Figure 4). In this dataset only splicing of the exon in ADD1, a known MBNL2 target (17), correlated significantly to MBNL2 expression. We found no further significant correlations, indicating that the degree of splicing regulation during brain development is much larger than the interindividual differences in splicing regulation in the adult brain.

### Prenatal-like splicing patterns in the DM1 brain are not driven by variation in mRNA expression of *CELF1* and *MBNL1/2*

We finally set out to investigate to what extent the coordinated splicing differences observed in the developing and the DM1 frontal cortex can be explained by variation in sample donor age and splicing factor RNA expression. For this analysis, we calculated pairwise correlations between the exon inclusion levels of all 34 high-confidence splice events. A pairwise correlation of two splice events captures the degree to which the two exons are both included or excluded (i.e. strong positive correlation) or are mutually exclusive (i.e. strong negative correlation). Next, we controlled the pairwise correlations for variation in age, splicing factor expression and disease state by calculating the partial correlations and compared the distribution of partial correlations to the zero-order (uncorrected) correlations (Figure 5). A difference between the partial and zero-order correlation suggests that the zero-order correlation can be partially explained by one of the factors that are controlled for in the partial correlation. In the context of the DM1 frontal cortex, we also considered the disease state (DM1 or unaffected).

**Figure 5:**
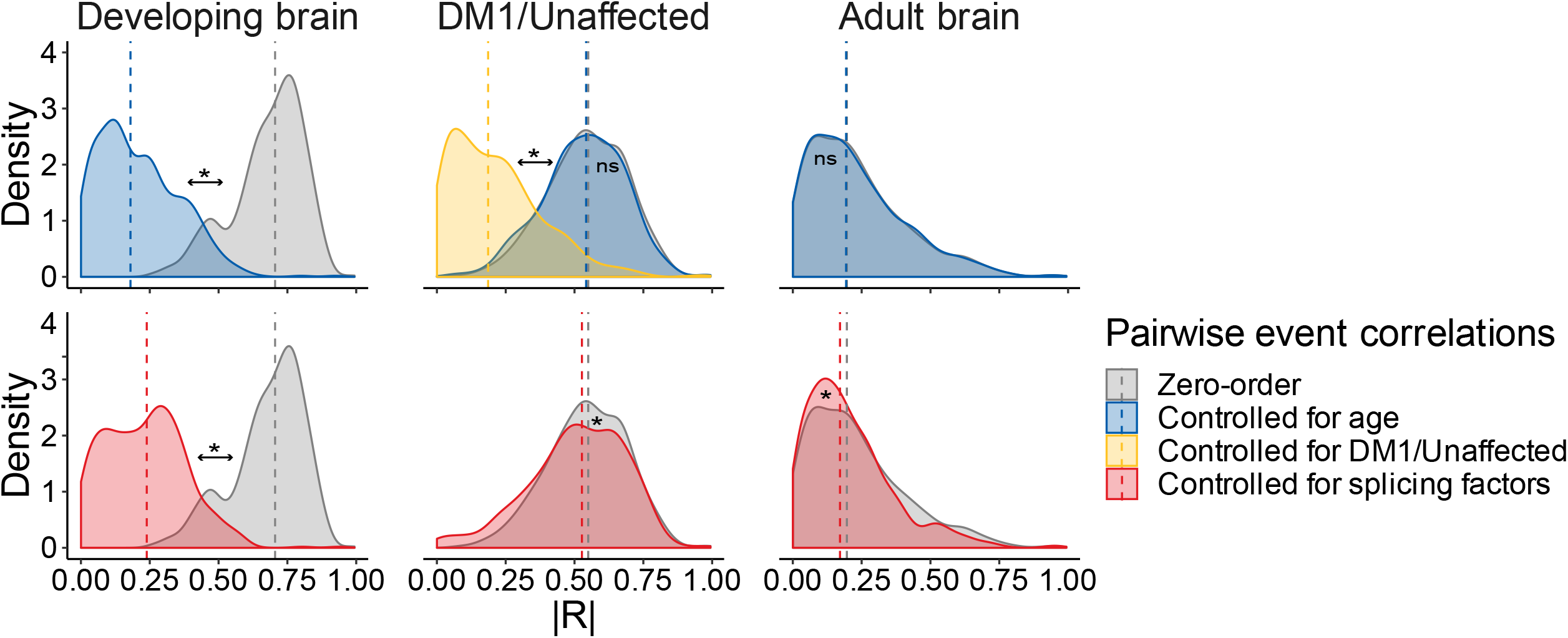
Pairwise correlations of splice events are partially explained by age differences and splicing factor RNA expression in the developing frontal cortex but not in the DM1 or unaffected adult frontal cortex. Pairwise (cross-) correlations were computed between the Ψ of all 34 high-confidence splice events. The density plots show the distribution of the absolute Spearman’s rank correlation coefficients for frontal cortex samples from the healthy, developing brain (left), from DM1 patients and unaffected adults (middle) and from the healthy, adult brain (right). Zero-order (i.e. uncorrected) pairwise correlations (in grey) were separately controlled for the following variables: age of the sample donor (in blue), disease state (DM1 or unaffected; in yellow) or RNA expression of *CELF1, MBNL1* and *MBNL2* (in red). Differences between zero-order distributions and their corrected counterparts were assessed by the rank-sum test. P-values were FDR-corrected with the Benjamini-Hochberg procedure (ns: p > 0.05,*: p < 0.05). The dashed vertical line indicates the median correlation.

As expected, controlling for age in the developing brain cohort, that features age as its main contrast, resulted in a strong decrease in pairwise correlations. Since the expression of CELF1 and MBNL1/2 was strongly related to the age of the subject, as shown in Figure 3, these factors could not be assessed independently and showed the same effect on the pairwise correlations. Nonetheless, the data support the notion that the coordinated changes in splicing during development are linked to a shift in the balance between CELF1 and MBNL1/2 expression. In the DM1 cohort, pairwise correlations were strongly decreased after controlling for disease state, but remained similar after controlling for age or splicing factor expression. Thus, we conclude that coordination of splicing in the DM1 frontal cortex is driven by differences in controls vs DM1 patients. In the healthy adult brain dataset pairwise correlations were generally low, and were unaltered by controlling for the factors mentioned above.

## DISCUSSION

In this work we offer a comprehensive overview of DM1 splice abnormalities in the frontal cortex that are related to development. We show that the majority of dysregulated splice events in frontal cortices in DM1 patients are changed towards prenatal splice variants, and identify 34 events with high confidence and a large shift in Ψ. This transcriptome-wide analysis provides insight in DM1 as a developmental disorder.

Various studies in humans and model organisms have shown that brain development goes hand-in-hand with changes in alternative splicing (14, 48). This was also observed in our analysis of the BrainSpan dataset, where we identified around 450 splice events that were different between prenatal and postnatal human frontal cortex. Of the 130 splice events that were dysregulated in the DM1 frontal cortex, 34 overlapped with the ∼450 developmentally regulated splice events, a significant enrichment. We focus on these 34 events because we assume that splice events that are developmentally regulated are more likely to have a physiological effect. Among these were known DM1-related splice events such as those in ADD1, CAMKK2, DMD, MAPT, MBNL1, MBNL2, and SORBS1 (21, 49), as well as several novel events. The splice events in GABRG2, NRCAM, ARHGAP44, DLGAP1 and CSNK1D were also found mis-spliced (|ΔΨ| ≥ 0.2, FDR ≤ 0.05) by Goodwin et al. in a previous RNA-Seq study on the DM1 frontal cortex, adding confidence to these exons for their relevance for DM1 (20).

Among the 34 events, altered splicing of GABRG2, TCF3, NRCAM, ARHGAP44, DLGAP, SEPT11, DCLK1 and CSNK1D may be most relevant for the DM1 brain phenotypes given their known functions in the CNS. Of these, only the events in SEPT11, DCLK1 and CSNK1D lead to a change in reading frame (exon length is not divisible by three), potentially leading to mRNA degradation or dysfunctional proteins. Still, e.g. for DCLK1 the identified splice event results in a different C-terminus of the protein, differentiating between DLCK1-alpha and -beta, of which it is known that alpha form is dominant in the prenatal brain (50). Of note, DCLK1 (isoform-) expression is associated to cognitive abilities (51) and anxiety (52).

Regarding CSNK1D (Casein kinase I isoform delta), exon 9 inclusion prevents further downstream translation of exon 10 by introducing a stop codon (53). Interestingly, this transcript variant alters the circadian rhythm and sleep alterations are a prevalent DM1 phenotype (54, 55). Ongoing efforts aim to develop small molecules for modulating CSNK1D activity but target specificity remains a major challenge (53).

GABRG2 (Gamma-Aminobutyric Acid Type A Receptor Subunit Gamma 2) has been associated with a plethora of neuro(developmental) disorders, mostly related to epilepsy (56–58). GABRG2 is known to be differentially spliced during development, where the short variant is expressed in early development (59). This alternatively spliced (micro-) exon is 24 nucleotides long and codes for an 8-amino acid stretch that alters GABA[A]R complex composition and modulates its activity (60). MS-based evidence has been gathered that, at least in rats, both isoforms are translated into protein (61). In schizophrenic patients the ratio of short (γ2S) vs. long (γ2L) GABRG2 isoforms is also altered, although in a way opposite to the DM1-situation presented in the data from Otero et al, with a relative increase of the short isoform (62). Additionally, GABRG2 variants have been observed in individuals affected by Dravet syndrome, autism spectrum disorder, developmental delay and intellectual disability (57, 63). Of major interest is that GABRG2 is a known drug target, and that the ratio of different isoforms can be altered via drugs (64–66).

Although not identified as dysregulated in the Goodwin study (20), TCF3 (Transcription Factor 3, or E2-alpha [E2A]) is a hit with a possibly important impact. TCF3 functions as a transcriptional regulator in neuronal differentiation, with an impact on many other genes (67). Interestingly, TCF3 can regulate IL-6 signaling, which is disturbed in DM1 (68, 69). Since there are differences in splicing aberrations between brain regions in DM1, our findings may not always be representative for other brain regions than the frontal cortex (70).

We analyzed the expression of the CELF and MBNL mRNAs throughout development and in the frontal cortex of adult DM1 patients and controls. In the literature, MBNL2 is generally considered the brain-related member of the MBNL family, but it is also known that MBNL1 can compensate for loss of MBNL2 function, or even plays an important role in the brain development itself (17, 71, 72). We noted that MBNL1, but not MBNL3, is also expressed in the human brain throughout development at levels comparable to those of MBNL2. This is confirmed via the Allen brain map and by recent single cell analyses (73, 74).

Consistent with data from the mouse heart (15), we observed a gradual increase in MBNL1/2 mRNA expression and a decrease of CELF1 expression with age in the human brain. The balance in MBNL1/2 and CELF activity during development is likely explained by these different transcription levels. This is especially relevant because many exons are sensitive to specific levels of MBNL1/2 activity (75). In turn, alterations in this balance are a likely cause of the regulation of a proportion of the developmentally regulated splicing events, as was demonstrated in the brains of Mbnl2 knock-out mice and muscles of Mbnl1 knock-out mice. In addition, there are exons in both MBNL1 (exon 5) and MBNL2 (exon 5 and 8) that are alternatively spliced throughout development (17, 76). These are likely part of an autoregulatory mechanism controlling the localization and activity of these splicing factors (9). MBNL1 exon 1 is also involved in MBNL1 autoregulation (9), but we found no differences in splicing in any of our analyses (data not shown).

In DM1 patients we observed a switch to fetal splice isoforms but this was not accompanied by a decrease but by a slight increase in the MBNL1/2 mRNA expression, while the CELF1 mRNA expression was unchanged. This confirms the basis of the altered balance between MBNL and CELF in DM1 patients: on the one hand MBNL loss-of-function is mainly caused by the differential splicing of MBNL1/2 and/or the entrapment of MBNL1/2 proteins in foci which is insufficiently compensated by an increase in MBNL1/2 transcription levels, and on the other hand the increased activity of CELF1/-2 is due to increased protein levels and/or their phosphorylation level (10, 77). In summary, MBNL1/2 and CELF1 seem transcriptionally regulated during development, but are heavily post-transcriptionally regulated in adult brains and muscles, e.g. through altered splicing and altered cellular localization (MBNL1/2) and post-translational modification (CELF1) (9, 78, 79).

In conclusion, we connected developmentally regulated splicing events with those dysregulated in DM1. This provides insights into the DM1 disease mechanism and helps to prioritize splice events for further investigation.

## ACKNOWLEDGMENT

The Brainspan datasets used for the analysis described in this manuscript were obtained from dbGaP at http://www.ncbi.nlm.nih.gov/gap through dbGaP accession number phs000731.v2.p1. Submission of the data, phs000731.v2.p1, to dbGaP was provided by Dr. Nenad Sestan. Collection of the data and analysis was supported by grants from the National Institutes of Health (MH089929, MH081896, and MH090047). Additional support was provided by the Kavli Foundation, a James S. McDonnell Foundation Scholar Award, NARSAD, and the Foster-Davis Foundation.

The Genotype-Tissue Expression (GTEx) Project was supported by the Common Fund of the Office of the Director of the National Institutes of Health (commonfund.nih.gov/GTEx). Additional funds were provided by the NCI, NHGRI, NHLBI, NIDA, NIMH, and NINDS. Donors were enrolled at Biospecimen Source Sites funded by NCI\Leidos Biomedical Research, Inc. subcontracts to the National Disease Research Interchange (10XS170), Roswell Park Cancer Institute (10XS171), and Science Care, Inc. (X10S172). The Laboratory, Data Analysis, and Coordinating Center (LDACC) was funded through a contract (HHSN268201000029C) to the The Broad Institute, Inc. Biorepository operations were funded through a Leidos Biomedical Research, Inc. subcontract to Van Andel Research Institute (10ST1035). Additional data repository and project management were provided by Leidos Biomedical Research, Inc. (HHSN261200800001E). The Brain Bank was supported supplements to University of Miami grant DA006227. Statistical Methods development grants were made to the University of Geneva (MH090941 & MH101814), the University of Chicago (MH090951, MH090937, MH101825, & MH101820), the University of North Carolina - Chapel Hill (MH090936), North Carolina State University (MH101819), Harvard University (MH090948), Stanford University (MH101782), Washington University (MH101810), and to the University of Pennsylvania (MH101822). The datasets used for the analyses described in this manuscript were obtained from dbGaP at http://www.ncbi.nlm.nih.gov/gap through dbGaP accession number phs000424.v8.p2.

---

**Supplemental Figure S1:**
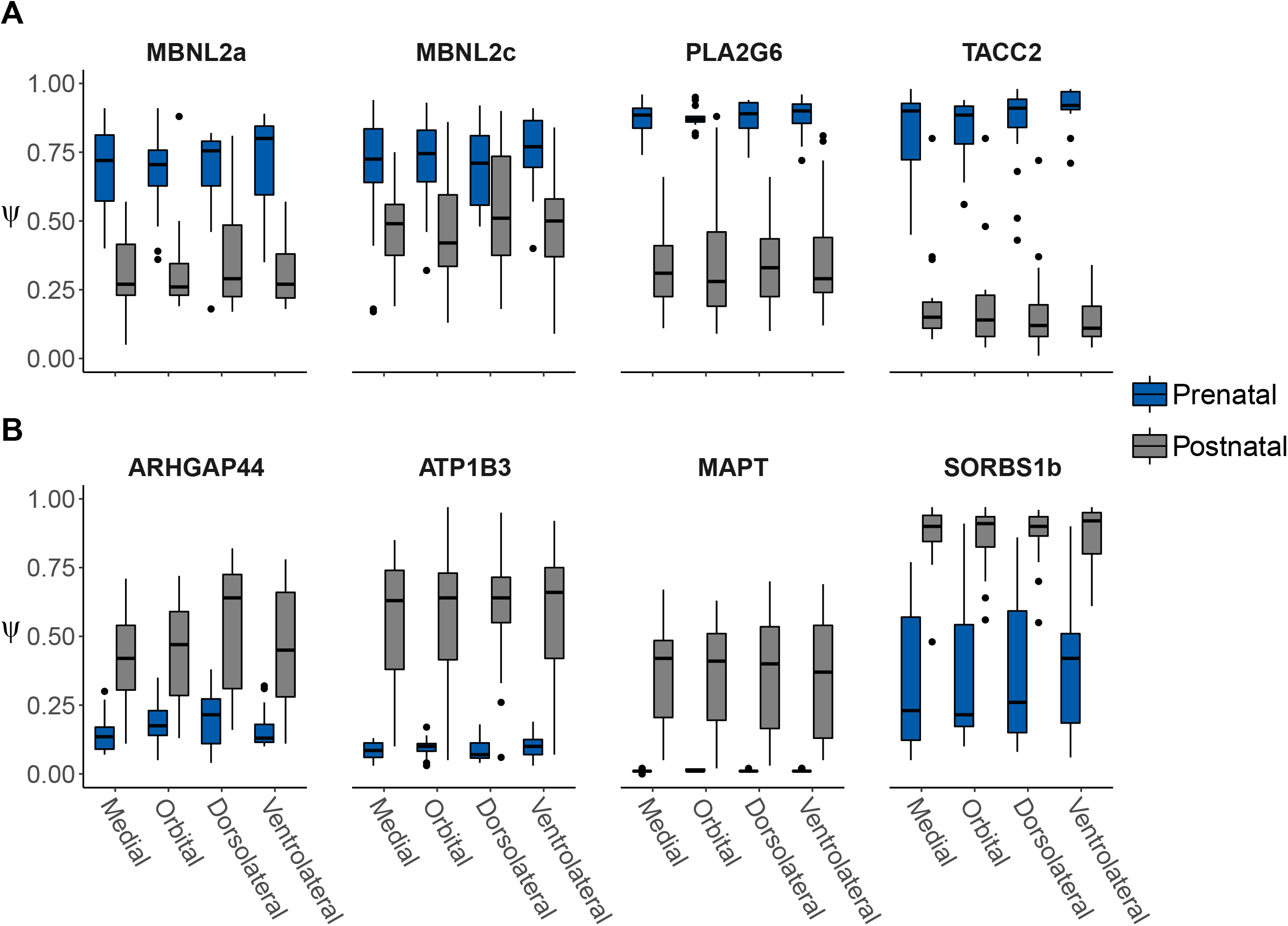
Exon inclusion does not differ significantly between subregions of the frontal cortex during human brain development. Differences in Ψ are shown for the selection of splice events in Figure 2. The boxplots are based on samples from frontal cortex subregions of DM1-unaffected prenatal and postnatal donors. A rank-sum test showed no significant difference between any subregions within the prenatal or postnatal group (p > 0.05). P-values were FDR-corrected with the Benjamini-Hochberg procedure.

**Supplemental Figure S2:**
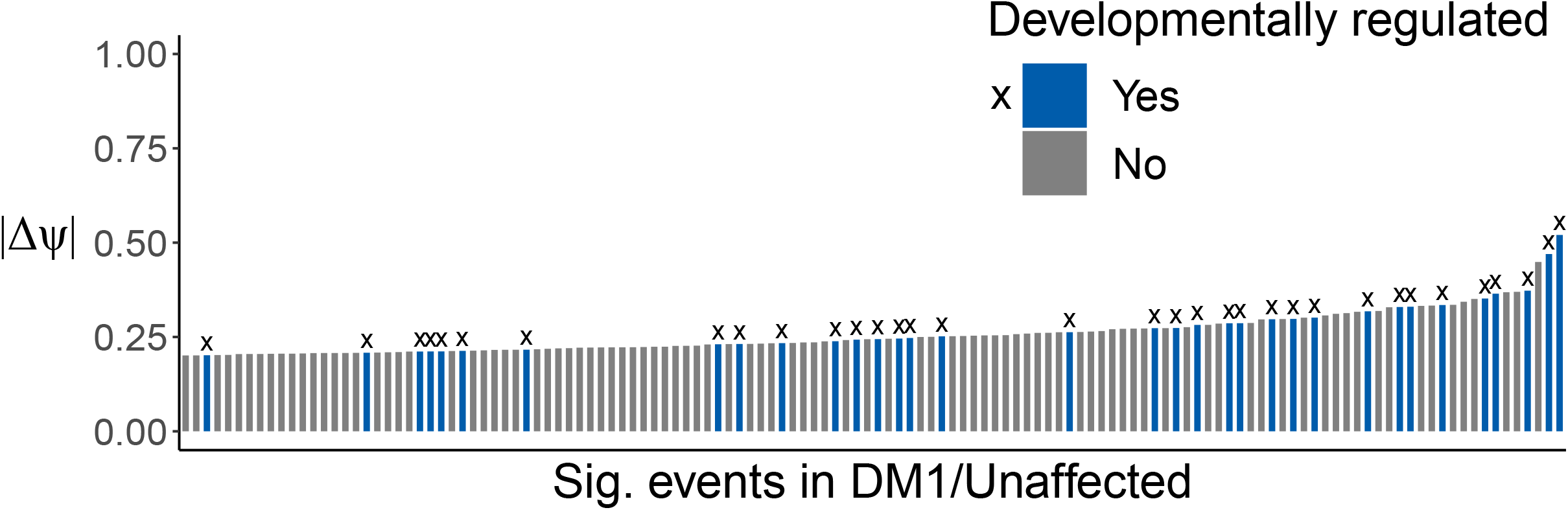
Developmentally-regulated splice events show on average larger differences in Ψ between DM1 patients and unaffected controls compared to all DM1-related events. The absolute Ψ difference is shown for 130 splice events that were found to be significantly different when comparing frontal cortex samples from DM1 patients and unaffected controls (|ΔΨ| > 0.2, p < 0.01 by rank-sum test). Splice events with a significant Ψ difference between prenatal and postnatal samples of the healthy, developing brain are colored in blue and marked by the letter “x”.

**Supplemental Figure S3:**
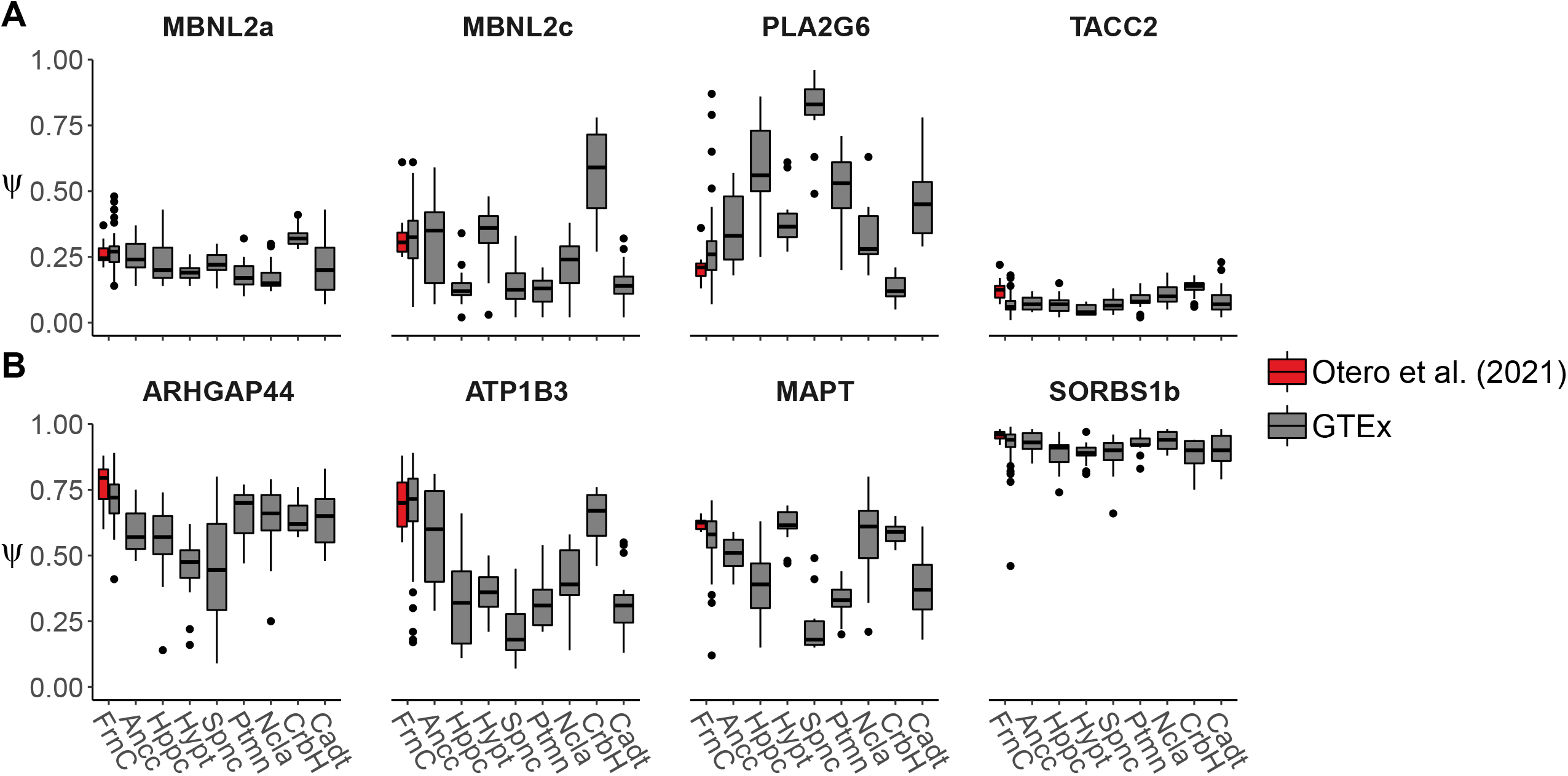
Exon inclusion varies between CNS regions in the healthy, adult brain. Differences in Ψ are shown for a selection of splice events with the largest decrease (A) or increase (B) in Ψ in DM1 patients compared to unaffected adults. The boxplots are based on samples from various CNS regions of unaffected, adult donors. No significant Ψ difference was found between control frontal cortex samples from Otero et al. (2021) and GTEx. The following regions are shown: FrnC=Frontal cortex (Otero et al., 2021: n=8; GTEx: n=54), Ancc=Anterior cingulate cortex (n=15), Hppc=Hippocampus (n=15), Hypt=Hypothalamus (n=14), Spnc=Spinal cord (n=14), Ptmn=Putamen (n=15), Ncla=Nucleus accumbens (n=15), CrbH=Cerebellar Hemisphere (n=15), Cadt=Caudate (n=15).

**Supplemental Figure S4:**
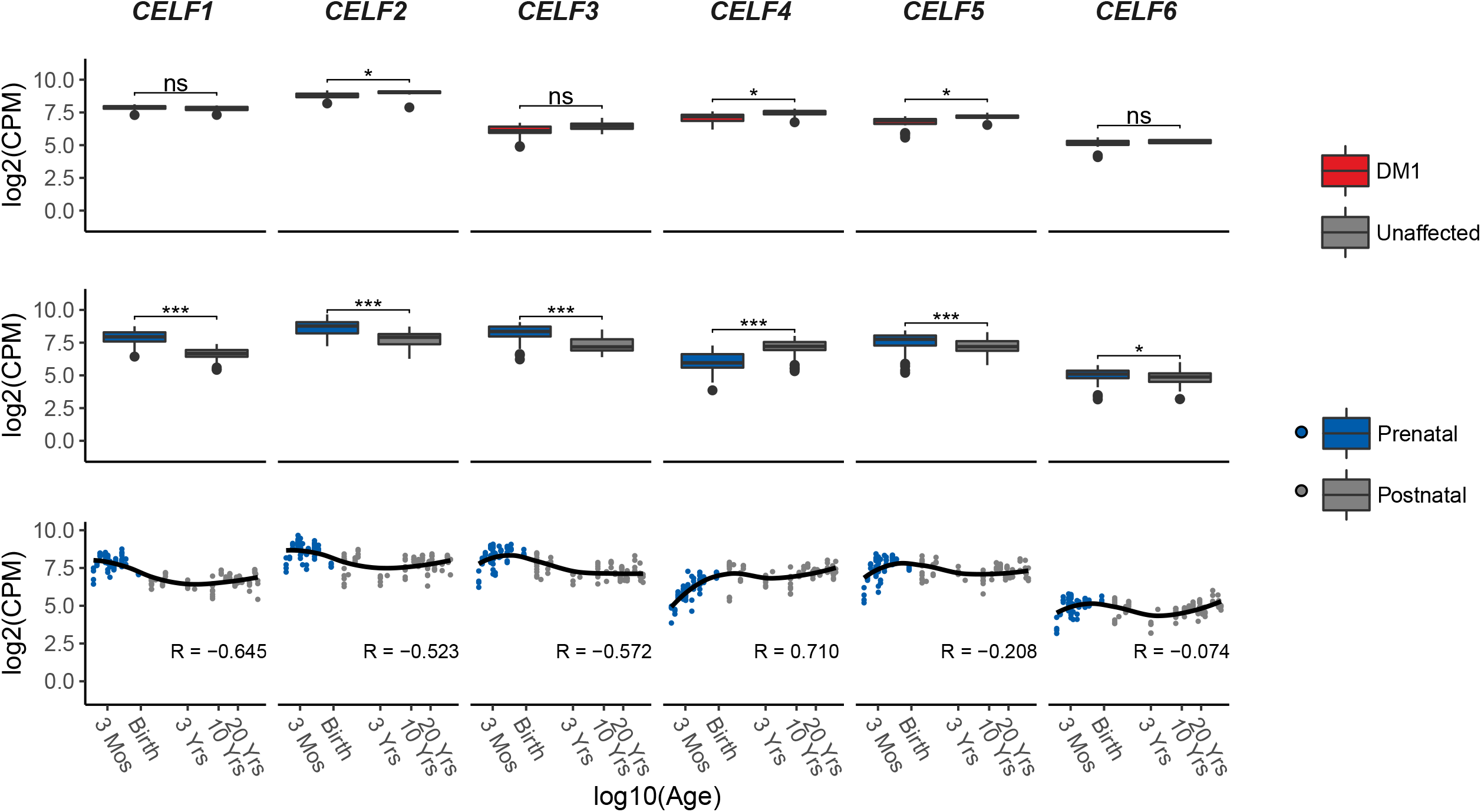
RNA expression of CELF family genes throughout human brain development and between DM1 patients and unaffected adults. Representation as in Figure 3.

**Supplemental Figure S5:**
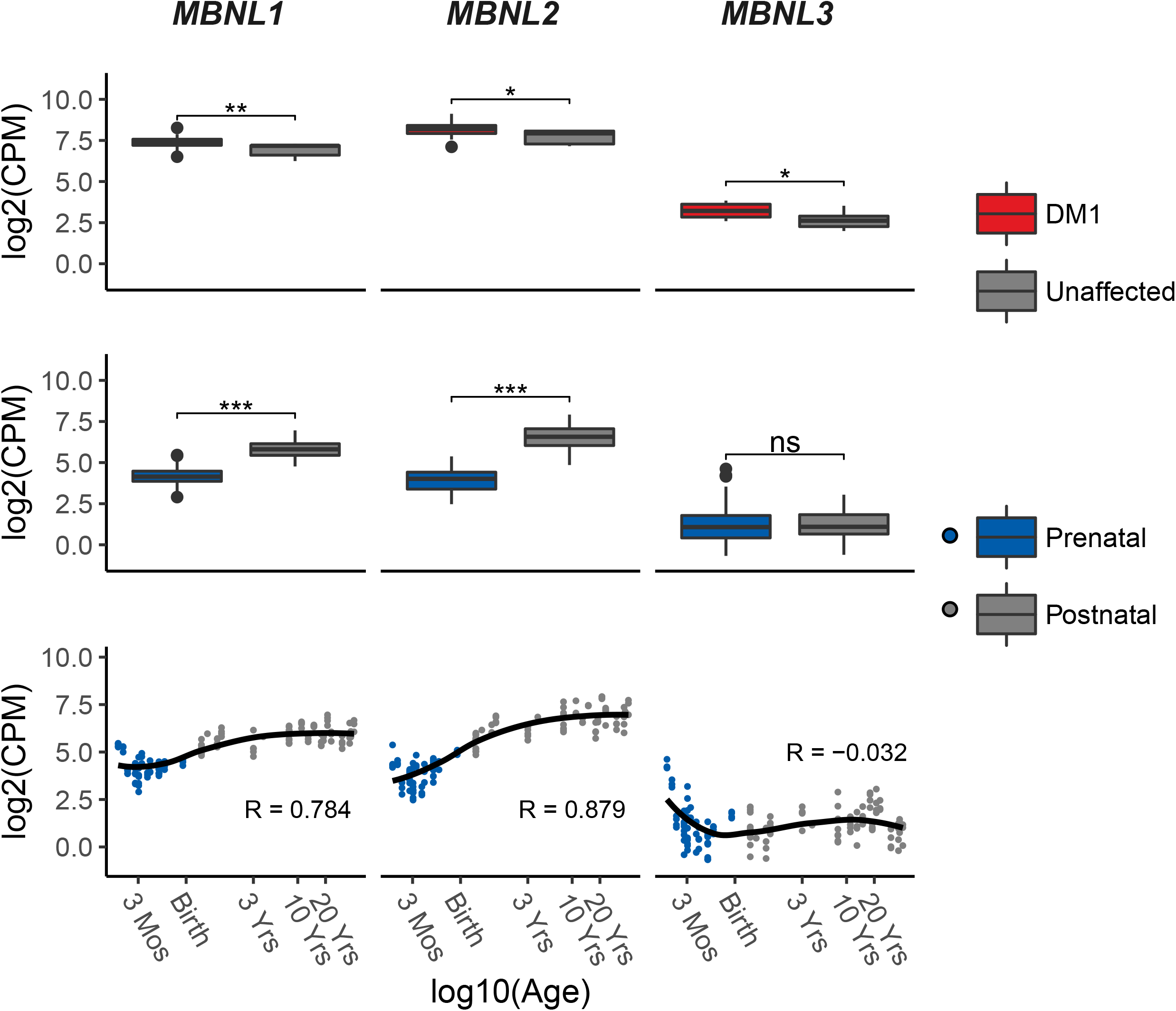
RNA expression of MBNL family genes throughout human brain development and between DM1 patients and unaffected adults. Representation as in Figure 3.

**Supplemental Figure S6:**
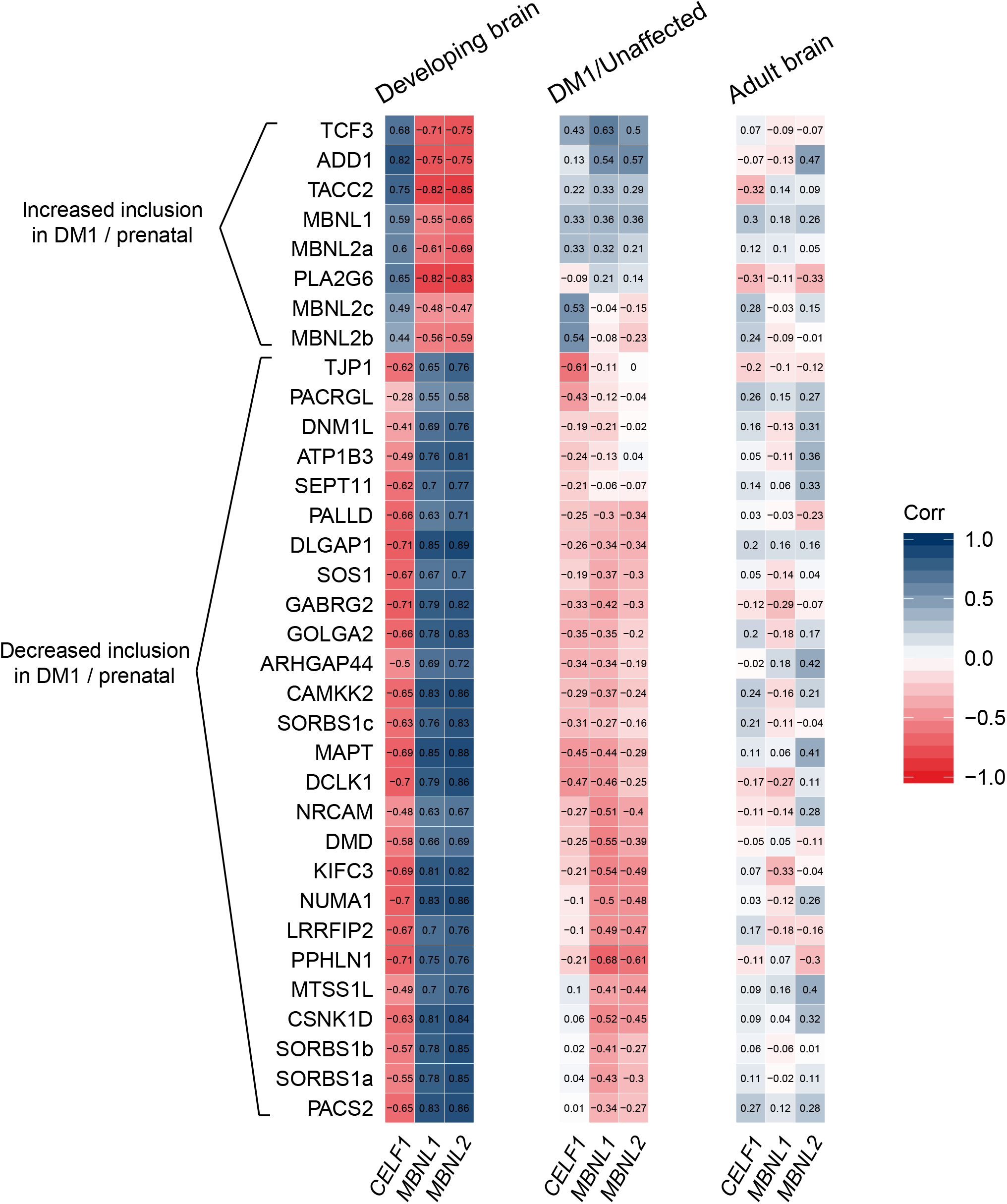
Correlation coefficients for correlation between high-confidence splice events and splicing factors expression. Representation as in Figure 4, supplemented with the value of the Spearman’s rank correlation coefficients.

**Supplemental Figure S7:**
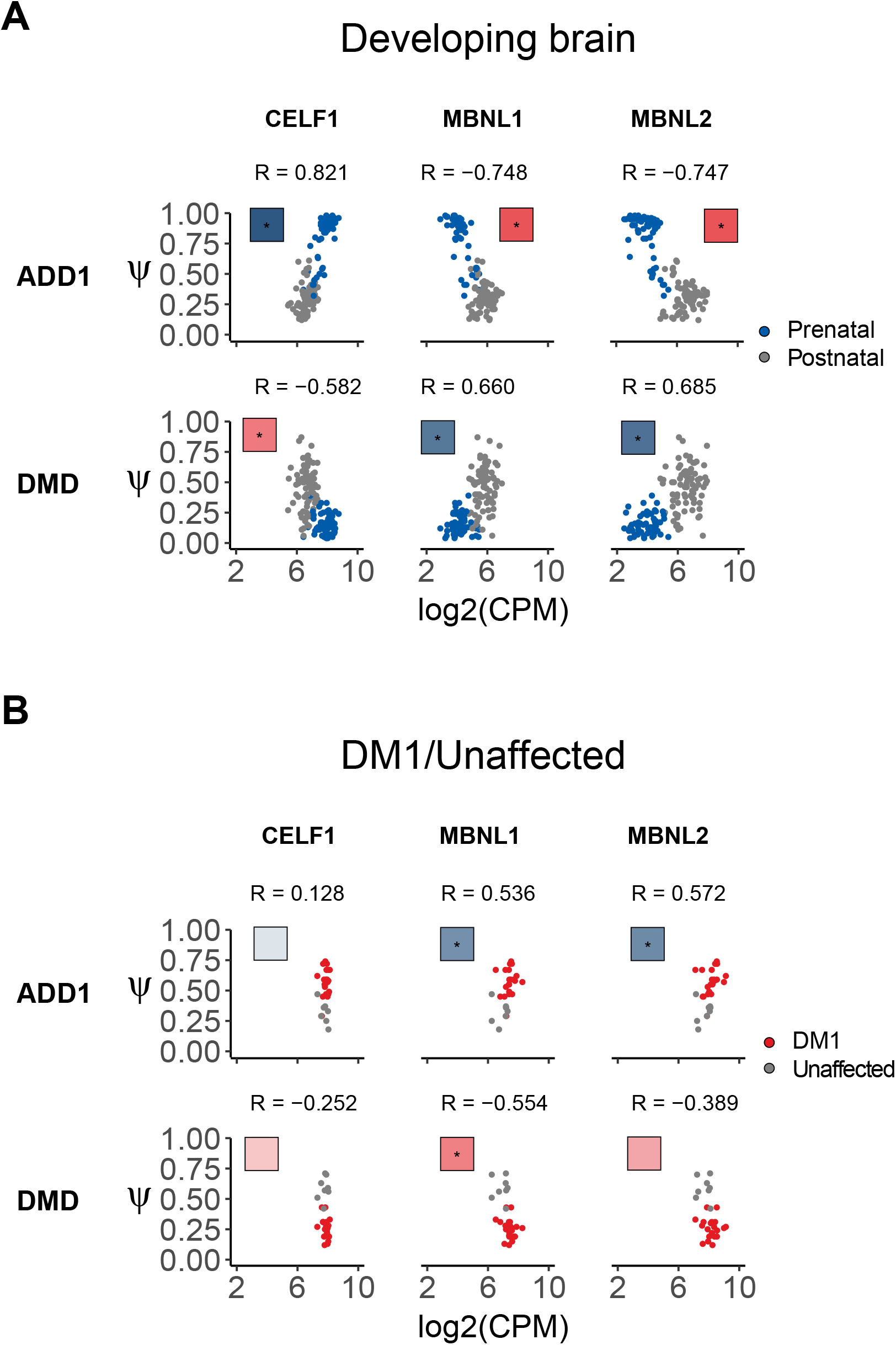
Scatter plots of ADD1 and DMD exon inclusion versus RNA expression of CELF1, MBNL1 and MBNL2. The Ψs of two exemplary splice events (i.e. ADD1 and DMD) are plotted against the RNA expression of *CELF1, MBNL1* and *MBNL2*. The scatter plots are based on frontal cortex samples from the healthy, developing brain (A) and from DM1 patients and unaffected controls (B). The saturation of the colored squares illustrates the strength of the correlation as in the heatmap in Figure 4. Asterisks indicate a significant correlation (FDR-corrected p < 0.05).

**Supplemental Figure S8:**
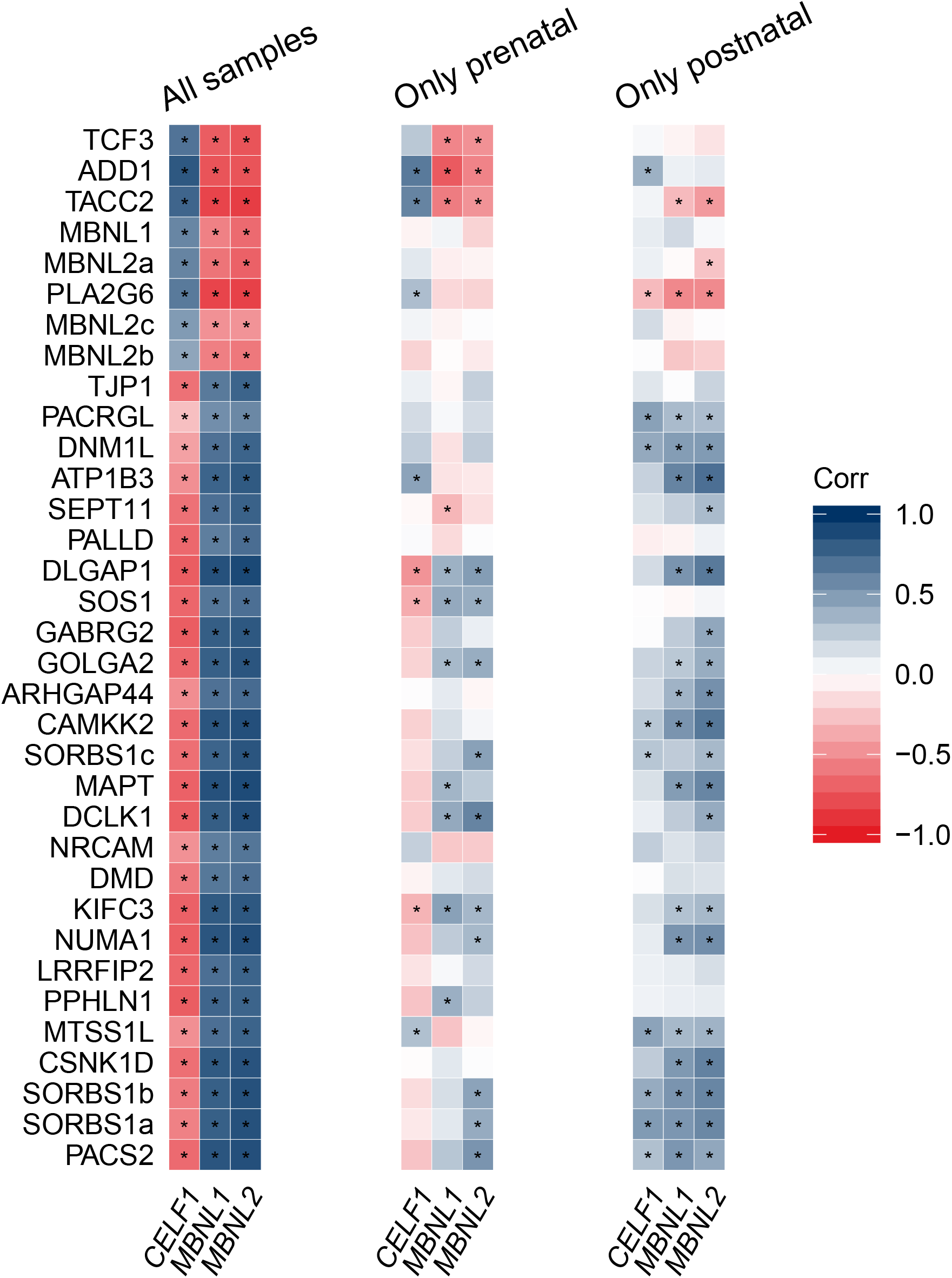
Mixed data from prenatal and postnatal sample donors is necessary to reveal all correlational patterns between the splice events. Correlation between the Ψs of high-confidence exon skipping events and RNA expression levels of *CELF1, MBNL1* and *MBNL2*. Correlations are displayed for frontal cortex samples from the healthy, developing brain before (left) and after (right) birth, along with their combination identical to the left panel in Figure 4 (left). Representation as in Figure 4.

**Supplemental Figure S9:**
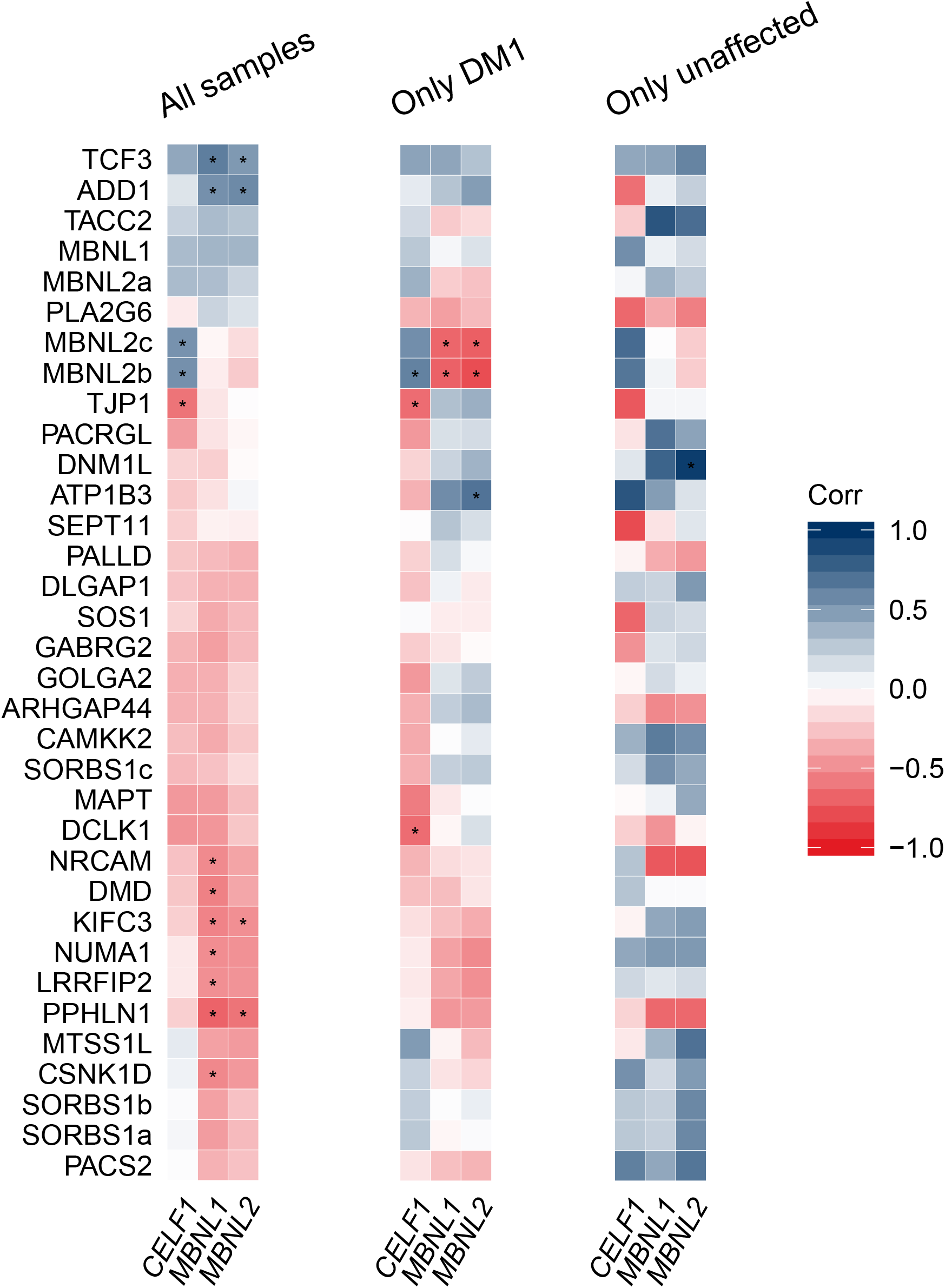
Mixed data from DM1 patients and unaffected adults is necessary to reveal weak correlational patterns between the splice events. Correlation between the Ψs of high-confidence exon skipping events and RNA expression levels of *CELF1, MBNL1* and *MBNL2*. Correlations are displayed separately for frontal cortex samples from DM1 patients (middle) and unaffected adults (right), along with their combination identical to the left panel in Figure 4 (left). Representation as in Figure 4.

